# *In vitro* efficacy of Artemisinin-based treatments against SARS-CoV-2

**DOI:** 10.1101/2020.10.05.326637

**Authors:** Kerry Gilmore, Yuyong Zhou, Santseharay Ramirez, Long V. Pham, Ulrik Fahnøe, Shan Feng, Anna Offersgaard, Jakob Trimpert, Jens Bukh, Klaus Osterrieder, Judith M. Gottwein, Peter H. Seeberger

**Affiliations:** Max Planck Institute for Colloids and Interfaces, Am Mühlenberg 1, 14476 Potsdam, Germany; Copenhagen Hepatitis C Program (CO-HEP), Department of Infectious Diseases, Hvidovre Hospital and Department of Immunology and Microbiology, Faculty of Health and Medical Sciences, University of Copenhagen, Copenhagen, Denmark; Freie Universität Berlin, Institute for Virology, Robert von Ostertag-Str. 7-13, 14163 Berlin, Germany; Department of Infectious Diseases and Public Health, Jockey Club College of Veterinary Medicine and Life Sciences, City University of Hong Kong, Kowloon Tong, Hong Kong; Freie Universität Berlin, Institute of Chemistry and Biochemistry, Arnimallee 22, 14195 Berlin, Germany

## Abstract

Effective and affordable treatments for patients suffering from coronavirus disease 2019 (COVID-19), caused by severe acute respiratory syndrome coronavirus 2 (SARS-CoV-2), are needed. We report *in vitro* efficacy of *Artemisia annua* extracts as well as artemisinin, artesunate, and artemether against SARS-CoV-2. The latter two are approved active pharmaceutical ingredients of anti-malarial drugs.

Proof-of-concept for prophylactic efficacy of the extracts was obtained using a plaque-reduction assay in VeroE6 cells. Subsequent concentration-response studies using a high-throughput antiviral assay, based on immunostaining of SARS-CoV-2 spike glycoprotein, revealed that pretreatment and treatment with extracts, artemisinin, and artesunate inhibited SARS-CoV-2 infection of VeroE6 cells. In treatment assays, artesunate (50% effective concentration (EC50): 7 μg/mL) was more potent than the tested plant extracts (128-260 μg/mL) or artemisinin (151 μg/mL) and artemether (>179 μg/mL), while generally EC50 in pretreatment assays were slightly higher. The selectivity index (SI), calculated based on treatment and cell viability assays, was highest for artemisinin (54), and roughly equal for the extracts (5-10), artesunate (6) and artemether (<7). Similar results were obtained in human hepatoma Huh7.5 cells. Peak plasma concentrations of artesunate exceeding EC50 values can be achieved. Clinical studies are required to further evaluate the utility of these compounds as COVID-19 treatment.

## Introduction

The pandemic with severe acute respiratory syndrome coronavirus 2 (SARS-CoV-2)^1, 2^ has worldwide been associated with over 1 million deaths from coronavirus disease 2019 (COVID-19).^3, 4, 5^ This febrile respiratory and systemic illness is highly contagious and in many cases life-threatening. Remdesivir, the only antiviral drug with proven *in vitro* and clinical efficacy, was approved for treatment of COVID-19.^6^ Still, COVID-19 treatment remains largely supportive with an urgent need to identify effective antivirals against SARS-CoV-2. An attractive approach is repurposing drugs already licensed for other diseases. Teas of *A. annua* plants have been employed to treat malaria in Traditional Chinese Medicine, as well as in clinical trials,^7, 8^ and are used widely in many African countries, albeit against WHO recommendations. Artemisinin (Figure 1, **1**), a sesquiterpene lactone with a peroxide moiety and one of many bioactive compounds present in *A. annua*, is the active ingredient to treat malaria infections.^9, 10^ The artemisinin derivatives artesunate (Figure 1, **2**) and artemether (Figure 1, **3**) exhibit improved pharmacokinetic properties and are the key active pharmaceutical ingredients (API) of WHO-recommended anti-malaria combination therapies used in millions of adults and children each year with few side effects.^11^ *A. annua* extracts are active against different viruses, including SARS-CoV.^12, 13, 14^ Therefore, we set out to determine whether *A. annua* extracts, as well as pure artemisinin, artesunate, and artemether are active against SARS-CoV-2 *in vitro*. Artemisinin-based drugs would be attractive repurposing candidates for treatment of COVID-19 considering their excellent safety profiles in humans, and since they are readily available for worldwide distribution at a relatively low cost.

**Figure 1.**
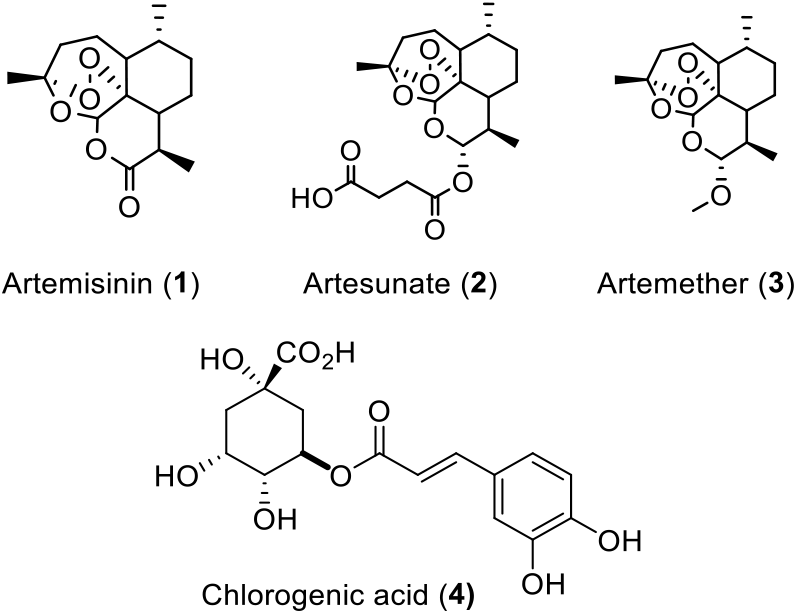
Artemisinin, related API derivatives artesunate and artemether, and chlorogenic acid.

## Results

### Extracts and compounds

*A. annua* plants grown from a cultivated seed line in Kentucky, USA, were extracted using either absolute ethanol or distilled water at 50 °C for 200 minutes, as described in Materials and Methods and Supplementary Information (Figure S1). For a third preparation, to protect artemisinin from degradation by reducing agents, ground coffee – a natural source of polyphenols such as chlorogenic acid (Figure 1, (**4**)) that also exhibit mild antiviral activities^15, 16^ – was coextracted with the plant material using ethanol (see supporting information). Solids were removed by filtration and the solvents were evaporated. The extracted materials were dissolved in dimethylsulfoxide (DMSO) (both ethanol extracts) or a DMSO:water mixture (3:1 for aqueous extract) and filtered (see supporting information for details). Artemisinin (Figure 1, (**1**)) was synthesized and purified following a published procedure,^17^ while artesunate (Figure 1, (**2**)) and artemether (Figure 1, (**3**)) were obtained from commercial sources.

### Plaque-reduction assays in VeroE6 cells for *in vitro* proof-of-concept of the pretreatment efficacy of *A. annua* extracts and artemisinin

To initially screen whether extracts and pure artemisinin were active against SARS-CoV-2, their antiviral activity was tested by pretreating VeroE6 cells at different time points during 120 minutes with selected concentrations of the extracts or compounds prior to infection with the first European SARS-CoV-2 isolated in München (SARS-CoV-2/human/Germany/BavPat 1/2020). The virus-drug mixture was then removed and cells were overlaid with medium containing 1.3% carboxymethylcellulose to prevent virus release into the medium. DMSO was used as a negative control. Plaque numbers were determined either by indirect immunofluorescence using a mixture of antibodies to SARS-CoV N protein^18^ or by staining with crystal violet.^19^ The addition of either ethanolic or aqueous *A. annua* extracts prior to virus addition resulted in reduced plaque formation in a concentration dependent manner with median effective concentration (EC50) values estimated to range between 5 and 168 μg/mL (Supplemental Figure S2A-C). Artemisinin exhibited little antiviral activity with an EC50 >220 μg/mL (Supplemental Figure S2D).

### Efficacy of *A. Annua* extracts in high-throughput antiviral *in vitro* assays in VeroE6 cells

Concentration-response experiments using the Danish SARS-CoV-2 isolate SARS-CoV-2/human/Denmark/DK-AHH1/2020 were performed employing a 96-well plate based high-throughput antiviral assay, allowing for multiple replicates per concentration, as described in Materials and Methods and Supplementary Information (Figures S3 and S4).^20^ Seven replicates were measured at each concentration and a range of concentrations was evaluated to increase data accuracy when compared to the plaque-reduction assay, which was carried out in duplicates. Extracts or compounds were added to VeroE6 cells either 1.5 h prior to (pretreatment (pt)) or 1 h post infection (treatment (t)), respectively, followed by a two-day incubation of virus with extracts or compounds. Both protocols yielded similar results, with slightly lower EC50 values observed for treatment assays.

The ethanolic extracts showed similar potency: for *A. annua* alone EC50 were 173 μg/mL (pt) and 142 μg/mL (t) and for *A. annua* with coffee EC50 were 176 μg/mL (pt) and 128 μg/mL (t) (Figures 2, 3 and Table 1). The aqueous extract was slightly less potent with EC50 being 390 μg/mL (pt) and 260 μg/mL (t) (Figures 2, 3 and Table 1). With all extracts, almost complete virus inhibition was achieved at high concentrations: For the *A. annua* ethanolic extract at 333 μg/mL (pt) and 444 μg/mL (t), for the *A. annua* + coffee ethanolic extract at 300 μg/mL (pt) and 267 μg/mL (t), and for the *A. annua* aqueous extract at 875 μg/mL (pt) and 1009 μg/mL (t) (Figures 2 and 3). The highest evaluated concentrations used in our assays were informed by the cytotoxicity of the extracts or compounds, as only concentrations resulting in cell viability greater than 90% were evaluated (Figures 2, 3, S5 and Table 1). Cell viability assays revealed median cytotoxic concentrations (CC50) of 1,044 μg/mL (*A. annua* ethanolic extract), 632 μg/mL (*A. annua* + coffee ethanolic extract), and 2,721 μg/mL (*A. annua* aqueous extract) (Figures 2, 3, S5 and Table 1). Selectivity indexes (SI) were determined by dividing CC50 by EC50 and revealed similar results for the *A. annua* ethanolic extract being 6 (pt) and 7 (t), the *A. annua* + coffee ethanolic extract being 3 (pt) and 5 (t) as well as the *A. annua* aqueous extract being 7 (pt) and 10 (t) (Table 1).

**Figure 2.**
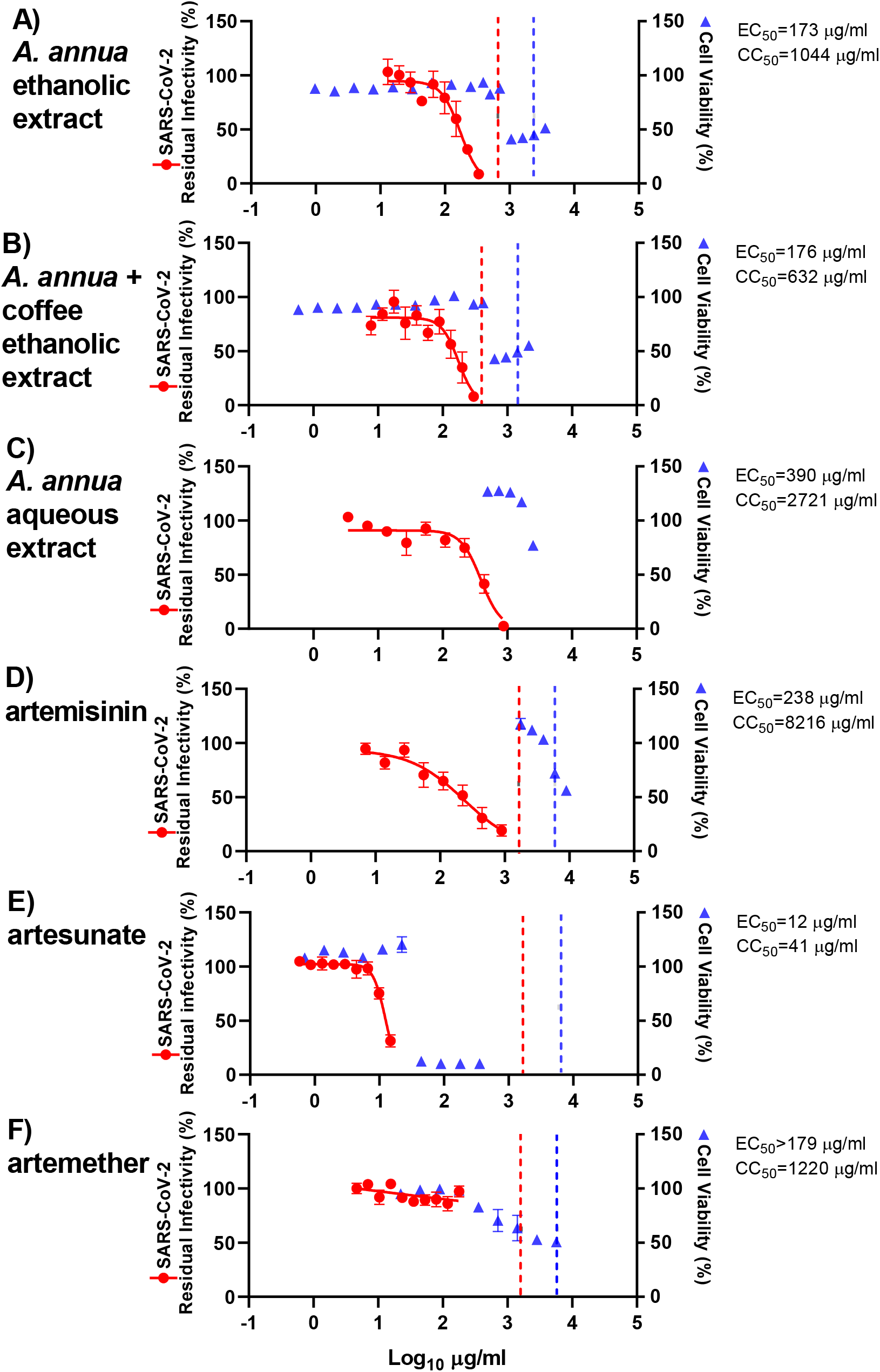
Pretreatment efficacy of extracts and compounds against SARS-CoV-2 in a high-throughput antiviral assay in VeroE6 cells. VeroE6 cells seeded the previous day in 96-well plates were treated with the specified concentrations of extracts (A) *A. annua* ethanolic extract, (B) *A. annua* + coffee ethanolic extract, and (C) *A. annua* aqueous extract, or compounds artemisinin (D), artesunate (E), and artemether (F) for 1.5 hours prior to infection with SARS-CoV-2. After a 2-day incubation, infected cells were visualized by immunostaining for SARS-CoV-2 spike glycoprotein and counted automatically as described in Materials and Methods. % residual infectivity for individual wells was calculated by relating counts of infected treated wells to the mean count of 14 infected nontreated control wells. Datapoints (red dots) are means of seven replicates with SEM. Sigmoidal dose response curves (red lines) were fitted and EC50 values were calculated in GraphPad Prism as described in Materials and Methods. % Cell viability and CC50 values were determined in replicate assays without infection with SARS-CoV-2 as described in Materials and Methods. Datapoints (blue triangles) are means of 3 replicates with SEM. The dotted red / blue lines indicate the concentrations at which an antiviral effect (<70% residual infectivity) / cytotoxic effect (<90% cell viability) due to DMSO is expected according to Figure S6.

**Table 1.** Efficacy of extracts and compounds in vitro.

| VeroE6 cells | EC50 (µg/ml)<sup>a</sup> |  | CC50 (µg/ml)<sup>b</sup> | SI<sup>c</sup> |
| --- | --- | --- | --- | --- |
|  | Pretreatment Assay | Treatment Assay |  |  |
| **Extract** |  |  |  |  |
| *A. annua* ethanolic extract | 173 | 142 | 1044 | 6 / 7 |
| *A. annua* + coffee ethanolic extract | 176 | 128 | 632 | 3 / 5 |
| *A. annua* aqueous extract | 390 | 260 | 2721 | 7 / 10 |
| **Compound** |  |  |  |  |
| Artemisinin | 238 | 151 | 8216 | 35 / 54 |
| Artesunate | 12 | 7 | 41 | 3 / 6 |
| Artemether | >179 | >179 | 1220 | <7 / <7 |
| **Huh7.5 cells** | EC50 (µg/ml)<sup>a</sup> |  | CC50 (µg/ml)<sup>b</sup> | SI<sup>c</sup> |
| **Extract** |  |  |  |  |
| *A. annua* ethanolic extract | 118 |  | 483 | 4 |
| **Compound** |  |  |  |  |
| Artemisinin | >208 |  | 5066 | <24 |
| Artesunate | 11 |  | 93 | 8 |
| Artemether | 135 |  | 303 | 2 |

<sup>a</sup> EC50, median effective concentration (µg/mL) was determined in VeroE6 cells in pretreatment or treatment antiviral assays or in Huh7.5 cells in treatment antiviral assays as described in Material and Methods. For artemether in VeroE6 cells and for artemisinin in Huh7.5 cells, <50% inhibition was observed at the highest non-cytotoxic concentration where cell viability was >90% of that of non-treated control cultures.
<sup>b</sup> CC50, median cytotoxic concentration (µg/mL) was determined as described in Material and Methods.
<sup>c</sup> SI, selectivity index, was determined as CC50 divided by EC50 based on results in pretreatment / treatment antiviral assays in VeroE6 cells or based on results in treatment antiviral assays in Huh 7.5 cells.

The two ethanolic extracts were diluted with DMSO that by itself caused reduction of cell viability to <90% when used at a 1:28 dilution, but not at dilutions ≥1:42 (Figure S6). Thus, the cytotoxicity observed when using the extracts at relatively high concentrations was most likely not caused by DMSO (Figures 2 and 3). DMSO at dilutions >1:152 including the dilutions used in antiviral assays did not have antiviral effects, defined as reduction of residual infectivity to <70% (Figure S6). Thus, the observed antiviral effect of the tested extracts was most likely not caused by DMSO. A pure coffee extract estimated to contain 2.5-fold higher coffee concentrations than the *A. annua* + coffee ethanolic extract did not result in reduction of cell viability to <90% at dilutions ≥1:28 (Figure S7). The cytotoxicity observed when using the *A. annua* + coffee extract at relatively high concentrations was most likely not caused by coffee (Figures 2 and 3). Interestingly, coffee extract alone showed some antiviral activity at dilutions ≤1:273 (Figure S7). Thus, the observed antiviral effect of the *A. annua* + coffee extract may be influenced by coffee.

**Figure 3.**
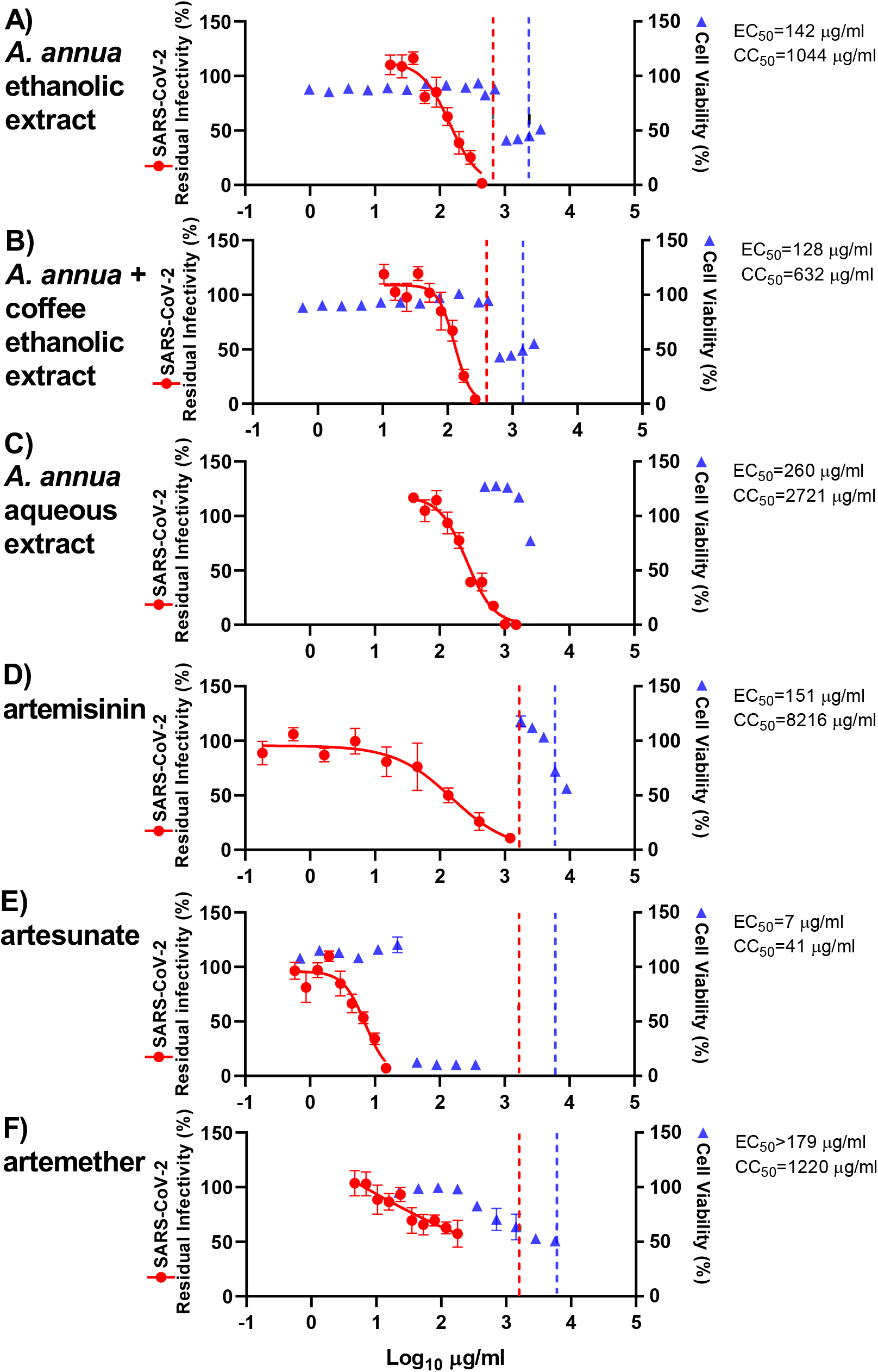
Treatment efficacy of extracts and compounds against SARS-CoV-2 in a high-throughput antiviral assay in VeroE6 cells. VeroE6 cells seeded the previous day in 96-well plates were infected with SARS-CoV-2 and after 1 hour incubation treated with the specified concentrations of extracts (A) *A. annua* ethanolic extract, (B) *A. annua* + coffee ethanolic extract, and (C) *A. annua* aqueous extract or compounds artemisinin (D), artesunate (E), and artemether (F). After a 2-day incubation, infected cells were visualized by immunostaining for SARS-CoV-2 spike glycoprotein and counted automatically as described in Materials and Methods. % residual infectivity for individual wells was calculated by relating counts of infected treated wells to the mean count of 14 infected nontreated control wells. Datapoints (red dots) are means of seven replicates with SEM. Sigmoidal dose response curves (red lines) were fitted and EC50 values were calculated in GraphPad Prism as described in Materials and Methods. % Cell viability and CC50 values were determined in replicate assays without infection with SARS-CoV-2 as described in Materials and Methods. Datapoints (blue triangles) are means of three replicates with SEM. The dotted red / blue lines indicate the concentrations at which an antiviral effect (<70% residual infectivity) / cytotoxic effect (<90% cell viability) due to DMSO is expected according to Figure S6.

### Efficacy of artemisinin and its derivatives in high-throughput antiviral *in vitro* assays in VeroE6 cell*s*

*A. annua* plants contain, in addition to many other bioactive compounds, artemisinin that is responsible for the potent anti-malarial activities of *A. annua*. To investigate whether artemisinin is the active component responsible for the antiviral activities of the plant extracts described above, the pure compound and synthetic derivatives were tested in pretreatment and treatment assays. Artemisinin was found to be active in SARS-CoV-2 assays with EC50 238 μg/mL (pt) and 151 μg/mL (t) (Figures 2, 3, and Table 1). Close to complete virus inhibition was achieved in both assays at the highest concentration evaluated in the assays, 893 (pt) and 1208 μg/mL (t). The SI for artemisinin is relatively high, 34 (pt) and 54 (t), based on a CC50 of 8,216 μg/mL (Figures 2, 3, S6, and Table 1). The observed cytotoxicity of artemisinin appeared to be at least partially caused by DMSO, as cytotoxicity was only observed at drug dilutions where DMSO was found to reduce cell viability (Figures 2, 3, and S6). The antiviral effects observed when using artemisinin at relatively high concentrations were most likely not due to the diluent DMSO (Figures 2, 3, and S6).

The synthetic artemisinin derivative artesunate, the API of WHO-recommended first-line malaria therapies with improved pharmacokinetic properties, showed the highest potency of all compounds tested, with EC50 being 12 μg/mL (pt) and 7 μg/mL (t) (Figures 2 and 3). In the treatment assay, close to complete virus inhibition was achieved at the highest evaluated concentration (15 μg/mL), as determined by cytotoxicity data, compared to 69% inhibition at this concentration in the pretreatment assay. Higher artesunate concentrations were not used considering its cytotoxicity in this assay (CC50: 41 μg/mL) (Figures 2, 3, S5, and Table 1). SI of 3 (pt) and 6 (t) were calculated (Table 1). The cytotoxicity and the antiviral effects observed when using artesunate at relatively high concentrations were most likely not due to the diluent DMSO (Figures 2, 3, and S6).

Artemether, another artemisinin-derivative that is used globally as the active ingredient in malaria medications, did not show a significant antiviral effect at concentrations of up to 179 μg/mL (Figures 2 and 3). Considering artemether’s cytotoxicity (CC50 of 1,220 μg/mL), an SI < 7 was calculated (Figures 2, 3, S5, and Table 1). The cytotoxicity observed when using artemether at relatively high concentrations was most likely not due to the diluent DMSO (Figures 2, 3, and S6).

### Efficacy of artemisinin-based treatment in high-throughput antiviral *in vitro* assays using Huh7.5 cells

The observed antiviral activity in these assays is effected by the ability of the pure compounds, and the compounds contained in the extracts, to enter the cells as well as their rates of metabolism within the cells. To exclude major differences in potency of extracts and compounds in human cells, treatment assays were also carried out in human hepatoma Huh7.5 cells, adding extracts or compounds to the cells immediately post infection. Overall, the ethanolic *A. annua* extract, artemisinin, artesunate, and artemether showed similar efficacy in Huh7.5 compared to VeroE6 cells. Artesunate (EC50: 11 μg/mL) was again found to be the most potent compound with close to complete virus inhibition at 22 μg/mL and an SI of 8 as determined by a CC50 of 93 μg/mL (Figures 4, S8 and Table 1). Artemether, (EC50: 135 μg/mL) with close to complete virus inhibition at 179 μg/mL, had an SI of only 2, based on CC50 of 303 μg/mL (Figures 4, S8 and Table 1). In Huh7.5 cells, the EC50 for the ethanolic *A. annua* extract was 118 μg/mL, with 76% virus inhibition at the highest evaluated concentration (150 μg/mL), as determined by cytotoxicity data; the CC50 was 483 μg/mL and the SI was 4 (Figures 4, S8 and Table 1). Artemisinin showed no significant virus inhibition at the highest evaluated concentration (208 μg/mL) and an SI <24, based on a CC50 of 5,066 μg/mL (Figures 4, S8 and Table 1).

**Figure 4.**
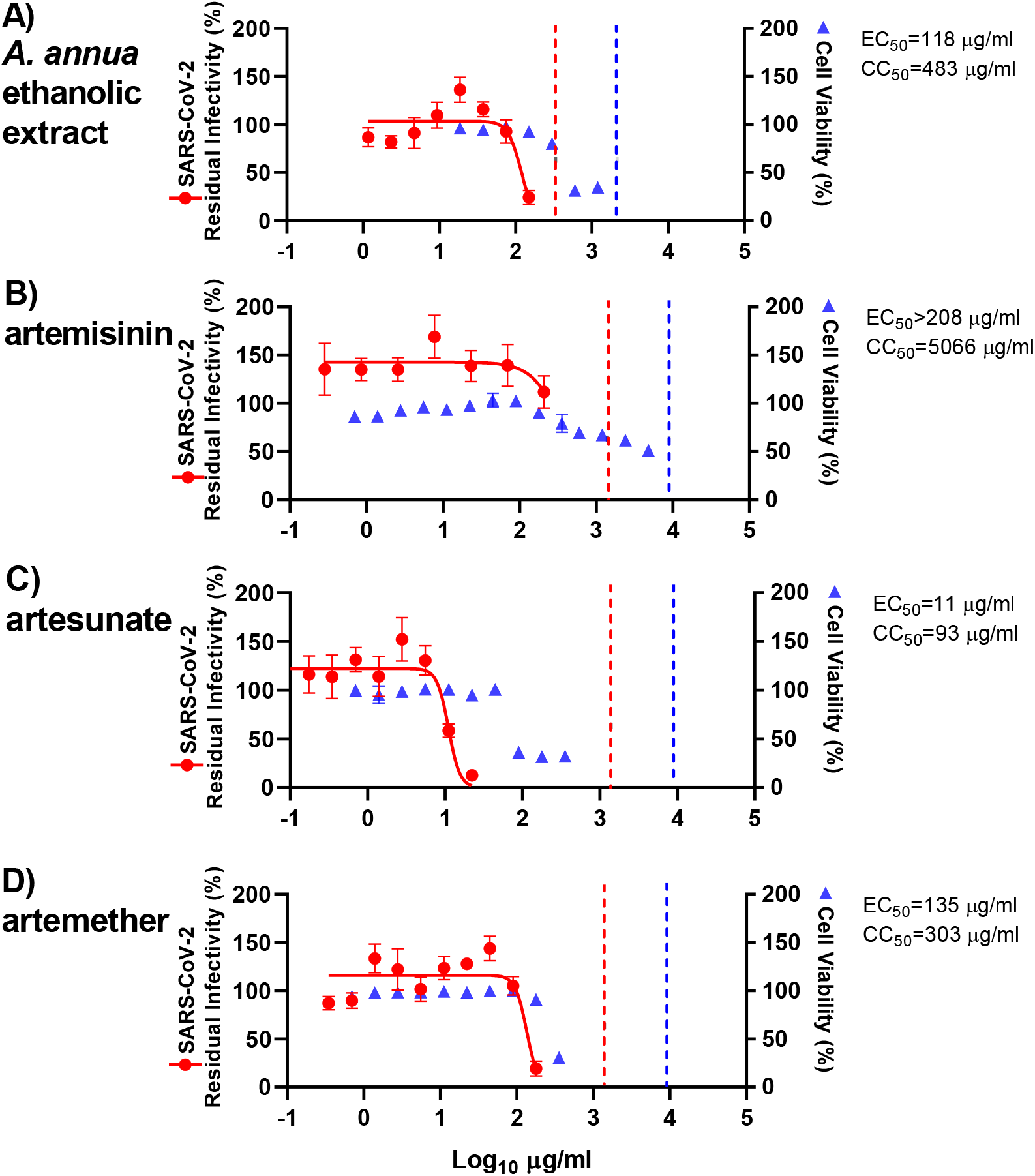
Treatment efficacy of extracts and compounds against SARS-CoV-2 in a high-throughput antiviral assay in Huh7.5 cells. Huh7.5 cells seeded the previous day in 96-well plates were infected with SARS-CoV-2 and directly treated with the specified concentrations of extract (A) *A. annua* ethanolic extract or compounds artemisinin (D), artesunate (E), and artemether (F). After a 3-day incubation, infected cells were visualized by immunostaining for SARS-CoV-2 spike glycoprotein and counted automatically as described in Materials and Methods. % residual infectivity for individual wells was calculated by relating counts of infected treated wells to the mean count of 14 infected nontreated control wells. Datapoints (red dots) are means of seven replicates with SEM. Sigmoidal dose response curves (red lines) were fitted and EC50 values were calculated in GraphPad Prism as described in Materials and Methods. % Cell viability and CC50 values were determined in replicate assays without infection with SARS-CoV-2 as described in Materials and Methods. Datapoints (blue triangles) are means of 3 replicates with SEM. The dotted red / blue lines indicate the concentrations at which an antiviral effect (<70% residual infectivity) / cytotoxic effect (<90% cell viability) due to DMSO is expected according to Figure S9.

In Huh7.5 cells, DMSO caused reduction of cell viability to <90% when used at a 1:28 dilution, but not at dilutions ≥1:56 (Figure S9). Thus, the cytotoxicity observed when using the ethanolic extract or the pure compounds at relatively high concentrations was most likely not caused by DMSO (Figure 4). DMSO at dilutions >1:179 including dilutions used in antiviral assays did not have any antiviral effects (Figure S9). Thus, the observed antiviral effect of the ethanolic *A. annua* extract and the pure compounds was most likely not caused by DMSO.

## Discussion

Here, we demonstrate the *in vitro* efficacy of artemisinin-based treatments against SARS-CoV-2. Initially, several *A. annua* extracts, as well as artemisinin, were screened for antiviral activity using a plaque-reduction assay in a pretreatment setting using a German SARS-CoV-2 strain from Munich. Based on these findings, three *A. annua* extracts and pure, synthetic artemisinin, artesunate, and artemether were studied in detail to establish concentration-response curves for extracts and compounds for pretreatment and treatment settings using a Danish SARS-CoV-2 strain from Copenhagen.

High-throughput antiviral assays facilitated testing of drug concentrations in multiple replicates resulting in accurate EC50 values. The EC50 values in the pretreatment setting were slightly higher than EC50 values determined in the treatment setting possibly because preincubation may have a negative impact on the stability of the extracts and pure compounds. Generally, EC50 values depend on the specific assay employed. While the type of assay we used with a single treatment and subsequent incubation of virus and drug is state of the art for antiviral efficacy measurements, assay modifications, such as repeated administration of treatment, might result in slightly different EC50 values. Since the active antiviral substance may be an artemisinin metabolite, such that the artemisinin derivatives and extracts can be considered prodrugs, we used the human Huh7.5 cell line to confirm the EC50 determined in VeroE6 cells.

While *A. annua* extracts have been considered “natural combination therapies” as they contain several bioactive compounds,^21^ the WHO discourages the use of non-pharmaceutical forms of artemisinin as a therapeutic option for malaria due to lack of standardization with its sourcing and preparation, implying risks of suboptimal efficacy and resistance development.^22^ In this context, it is important to note that the extracts used in this study were prepared from plants grown under optimized and standardized conditions, in a manner where concentrations of the extracted material are reproducible.

Interestingly, we found that coffee extracts exhibited *in vitro* efficacy against SARS-CoV-2. While modelling studies suggested that ingredients in coffee such as chlorogenic acid, caffeic acid, and tannins show activity against SARS-CoV-2,^23^ we here provide *in vitro* evidence of such an effect. Future studies are needed to elucidate the effect of coffee and active ingredients on SARS-CoV-2 in detail. In addition, future studies might address whether *A. annua* or active ingredients and coffee extracts show antagonistic, additive, or synergistic effects on SARS-CoV-2.

While compiling data for this study, Cao et al. reported efficacy of artemisinin derivatives against a SARS-CoV-2 isolate from Wuhan in VeroE6 cells.^24^ While extracts were not studied, the efficacy of artesunate was similar with EC50 of 13 μM compared to 18 μM in our VeroE6 treatment assay. Interestingly, EC50 for artemether and artemisinin were 8 fold and >8 fold higher in our VeroE6 treatment assay compared to values reported by Cao *et al*. This difference might be due to the nature of the assay employed. The assay used by Cao *et al*. was based on viral RNA determinations, the viral inoculum was removed post infection / prior to treatment and perhaps most importantly, the assay was terminated 24 hours post infection, which is expected to result in comparatively lower EC50 for compounds with an immediate antiviral effect but a limited capacity to control the virus, either due to limited antiviral efficacy or due to limited stability during a 48-hour treatment period. Importantly, we report for the first time the efficacy of the compounds in a human cell line (Huh7.5) in addition to efficacy in the monkey cell line VeroE6. Finally, in our study we confirm efficacy of artemisinin-based treatment for two European SARS-CoV-2 strains from Germany and Denmark, which are more closely related to the majority of SARS-CoV-2 strains circulating worldwide than the Wuhan strain.

Artesunate, the API in Food and Drug Administration (FDA) approved malaria treatments, showed the highest potency against SARS-CoV-2 among the extracts and pure compounds tested in VeroE6 and Huh7.5 cells. The extracts proved more potent than artemisinin and artemether that exhibited only borderline antiviral efficacy considering results in VeroE6 and Huh7.5 cells. With SI of below 10, except for artemisinin, a relatively low therapeutic window exists. It should be noted that certain drugs such as digoxin with SI values as low as 2 are used successfully in the clinic.^25^ Among the tested extracts and pure compounds, only artesunate showed EC50 values in the range of clinically achievable plasma and tissue concentrations. When the typically used doses of 2 to 2.4 mg/kg intravenously were administered, reported peak plasma concentrations (Cmax) were between 19.4 and 29.7 μg/mL in patients.^26^ Based on these observations and our treatment data in VeroE6 and Huh7.5 cells, the calculated Cmax/EC50 values are between 2.5 and 4.2. In animal studies following administration of a single dose of artesunate, tissue concentrations including lung, kidney, intestine, and spleen concentrations were several-fold higher than plasma concentrations.^27^ In contrast, following administration of artemisinin, artemether, and *A. annua* teas, Cmax values between 311-776 ng/mL were reported, which is close to three orders of magnitude below EC50 values for SARS-CoV-2. Plasma and tissue concentrations that can be achieved with standardized *A. annua* extracts with high artemisinin content used in this study still have to be determined. *In vivo*, immunomodulatory effects of artemisinin-based treatments have been reported for this class of drugs.^28^ Such effects that may involve cytokine signaling cannot be monitored in *in vitro* assays performed here and will have to be carefully studied in subsequent clinical evaluations.

## Materials and Methods

### Extraction

Solvent (250 mL ethanol or distilled water) was heated to 50 °C in an Erlenmeyer flask. Dried plant material (50g for ethanol, 25g for water) or dried plant material and preground coffee (50g, 50g) was added to the solvent and stirred for 200 minutes. The mixture was filtered and solid material washed with fresh ethanol or water. The solvent was removed by rotary evaporation and solid material stored at −30 °C prior to sample preparation.

### Sample Preparation

Dried extract was warmed to room temperature. The required sample mass was removed using a spatula. DMSO (3 mL, ethanol extracts) or DMSO:water (3:1, 8 mL water extract) was added and the mixture was heated (40 °C) to ensure solvation. The solution was filtered using a syringe filter and stored in a snap-close vial.

### Cell Culture

At FU Berlin, african green monkey kidney VeroE6 cells (ATCC CRL-1586) were maintained at 37 °C with 5% CO_2_ in Minimum Essential Medium (MEM; PAN Biotech, Aidenbach, Germany) supplemented with 10% fetal bovine serum (PAN Biotech), 100 IU/mL penicillin G and 100 μg/mL streptomycin (Carl Roth, Karlsruhe, Germany).

At CO-HEP, african green monkey kidney VeroE6 cells (kind gift from J. Dubuisson) as well as human hepatoma Huh7.5 cells^28^ were maintained at 37 °C with 5% CO_2_ in Dulbecco’s Modified Medium (DMEM) (Invitrogen, Paisley, UK) containing 10% heat inactivated fetal bovine serum (FBS) (Sigma, Saint Louis, Missouri, USA) and 100 U/mL penicillin + 100 μL streptomycin (Gibco/Invitrogen Corporation, Carlsbad, California, USA). Cells were subcultured every 2-3 days using trypsin (Sigma, Saint Louis, Missouri, USA) to maintain a subconfluent cell layer.

### Virus isolates

The SARS-CoV-2 BavPat 1 isolate (SARS-CoV-2/human/Germany/BavPat 1/ 2020 was provided by Dr. Daniela Niemeyer and Dr. Christian Drosten (Charité, Berlin, Germany) and obtained from an outbreak in Munich, Germany, in February 2020 (BetaCoV/Germany/BavPat1/2020).

The SARS-CoV-2/human/Denmark/DK-AHH1/2020 virus for cell culture studies was obtained following inoculation of VeroE6 cells with patient swab sample, virus propagation in VeroE6 cells and generation of a sequence confirmed 2nd viral passage stock with an infectivity titer of 5.5 log TCID50/mL as described in Ramirez *et al*.^20^

### Plaque reduction antiviral assay

Antiviral activity of artemisinin derivatives was evaluated on VeroE6 cells grown overnight in 12-well plates (Sarstedt) at a density of approximately 5×10^5^ cells/well. Cells were incubated in the presence of ten-fold serial dilutions of the compounds for 15 min, 30 min, 60 min or 120 min, before the virus was added at a concentration of approximately 200 plaque-forming-units (PFU) per well for 120 min. The virus-drug mixture was removed, and cells were overlaid with MEM-FBS containing 1.3% carboxymethylcellulose to prevent virus release into the medium. DMSO in cell culture medium at a 1:100 dilution (the highest concentration relative to the preparations of extracts / compounds) was used as a negative control, and virus plaque numbers were determined by manual counting of plaques following indirect immunofluorescence (IF) using a mixture of antibodies to SARS-CoV N protein^18^ or following staining with crystal violet.^19^ For IF, cells were fixed with 4% formalin and permeabilized with 0.25% Triton X-100. Unspecific binding was blocked with 1% FBS in phosphate buffered saline (PBS) containing 0.25% Triton X-100 (PBS-T) at room temperature for 30 min. Cells were incubated with the anti-N monoclonal antibodies (1:25 dilution in PBS-T) for 45 min, followed by incubation with secondary antibody (Alexa 488-labeled goat anti-mouse at a 1:500 dilution; Thermo Fisher). In each assay, each concentration was tested in one replicate culture; 5 infected and DMSO control treated cultures were included in each assay. Plaque counts recorded in each infected treated culture were related to the average count of the five control cultures to calculate the number of plaques as percent relative to the control. Two independent assays were carried out. Datapoints are means of two replicate cultures from the two independent assays with error bars reflecting the standard deviations (SD) (Figure S2). Selected concentrations were only tested in one of the assays and for these datapoints are based on single replicates. The MOI for infection was chosen aiming at on average 150-250 plaques per culture.

### High-throughput pretreatment and treatment antiviral assay in VeroE6 cells

96-well based antiviral assays in VeroE6 cells were developed based on assays previously established for evaluation of the efficacy of antivirals against hepatitis C virus.^30, 31^ VeroE6 cells were plated at 10,000 cells per well of poly-D-lysine-coated 96-well plates (Thermo Fisher Scientific, Rochester, NY, USA). For pretreatment assays, the next day, medium was exchanged to medium containing extracts or compounds adding 50 μL per well. After 1.5 h of incubation at 37 °C and 5% CO_2_, cells were inoculated with SARS-CoV-2/human/Denmark/DK-AHH1/2020 at MOI 0.0016 by adding 50 μL of diluted virus stock per well, resulting in the specified concentrations of extracts or compounds. For treatment assays, the next day, medium was exchanged by adding 50 μL of fresh medium per well. Then, cells were inoculated with SARS-CoV-2/human/Denmark/DK-AHH1/2020 at MOI 0.0016 by adding 50 μL of diluted virus stock per well. After 1 hour of incubation at 37 °C with 5% CO_2_, 50 μL of medium containing extracts or compounds were added resulting in the specified concentrations; alternatively, 50 μL of medium containing diluent (DMSO) or additive (coffee extract) were added resulting in the specified dilutions. For both assays, each concentration/dilution was tested in seven replicates; 14 infected and nontreated as well as 12 noninfected and nontreated control wells were included in each assay. After 48±2 hours incubation at 37 °C and 5% CO_2_, cultures were immunostained for SARS-CoV-2 spike glycoprotein and evaluated as described below.

### High-throughput treatment antiviral assay in Huh7.5 cells

Huh7.5 cells were plated at 8,000 cells per well of flat bottom 96-well plates (Thermo Fisher Scientific, Roskilde, Denmark). The next day, cells were inoculated with SARS-CoV-2/human/Denmark/DK-AHH1/2020 at MOI 0.0198 by adding 50 μL of diluted virus stock per well. Directly after, 50 μL of medium containing extracts or compounds were added resulting in the specified concentrations; alternatively, 50 μL of medium containing diluent (DMSO) were added resulting in the specified dilutions. Each concentration was tested in seven replicates; 14 infected and nontreated as well as 12 noninfected and nontreated control wells were included in the assay. After 72±2 hours incubation at 37 °C and 5% CO_2_, cultures were immunostained for SARS-CoV-2 spike glycoprotein and evaluated as described below.

### Immunostaining and evaluation of 96-well plates for high-throughput antiviral assays

Cells were fixed and virus was inactivated by immersion of plates in methanol (J.T.Baker, Gliwice, Poland) for 20 min. Unless specified, immunostaining was done at room temperature. Plates were washed twice with PBS (Sigma, Gillingham, UK) containing 0.1% Tween-20 (Sigma, Saint Louis, Missouri, USA). Endogenous peroxidase activity was blocked by incubation with 3% H2O2 for ten minutes followed by two washes with PBS containing 0.1% Tween-20 and blocking with PBS containing 1% bovine serum albumin (Roche, Mannheim, Germany) and 0.2% skim milk powder (Easis, Aarhus, Denmark) for 30 minutes. Following removal of blocking solution, plates were incubated with primary antibody SARS-CoV-2 spike chimeric monoclonal antibody (Sino Biological #40150-D004, Beijing, China) diluted 1:5000 in PBS containing 1% bovine serum albumin and 0.2% skim milk powder overnight at 4 °C. Following two washes with PBS containing 0.1% Tween-20, plates were incubated with secondary antibody F(ab’)2-Goat anti-Human IgG Fc Cross-Adsorbed Secondary Antibody, HRP (Invitrogen #A24476, Carlsbad, CA, USA) or Goat F(ab’)2 Anti-Human IgG - Fc (HRP), pre-adsorbed (Abcamab#98595, Cambridge, UK) diluted 1:2000 in PBS containing 1% bovine serum albumin and 0.2% skim milk powder for 2 h. Following two washes with PBS containing 0.1% Tween-20, SARS-CoV-2 spike glycoprotein was visualized using DAB substrate (Immunologic # BS04-110, Duiven, Netherlands). Spike protein positive cells were counted automatically using an ImmunoSpot series 5 UV analyzer (CTL Europe GmbH, Bonn, Germany) as described.^30,31,32^ The average count of 12 noninfected nontreated control wells, which was usually <50, was subtracted from the count of each infected well. Counts recorded in each infected treated well were related to the average count of 14 infected nontreated control wells to calculate % residual infectivity. Datapoints are means of seven replicates with standard errors of the means (SEM). Sigmoidal dose response curves were fitted and EC50 values were calculated with GraphPad Prism 8.0.0 using a bottom constraint of 0 and the formula Y= Top/(1+10^((LogEC50-X)*HillSlope)). The MOI for infection was chosen aiming at on average 3000-4000 counts per well for VeroE6 cells and on average 300-600 counts per well for the less permissive Huh7.5 cells in infected nontreated control wells upon termination of the respective assays. Representative 96-well images from assays in VeroE6 cells are shown in Figure S3 and representative images of single wells are show in Figure S4.

### Cell viability assays in VeroE6 and Huh7.5 cells

To evaluate cytotoxic effects of the tested extracts, compounds, diluents (DMSO) and additive (coffee extract), cell viability was monitored using the CellTiter 96^®^ AQueous One Solution Cell Proliferation Assay (Promega, Madison, WI, USA). VeroE6 cells or Huh7.5 cells were plated at 10,000 or 8,000 cells per well of flat bottom 96-well plates, respectively (Thermo Fisher Scientific, Roskilde, Denmark). The next day, medium was exchanged to contain specified concentrations of extracts or compounds or dilutions of DMSO and coffee extract adding 100 μL per well. Each concentration or dilution was tested in 3 replicates; at least 6 nontreated control wells were included in the assay. For VeroE6 cells, after 48±2 h, and for Huh7.5 cells, after 72±2 h of incubation at 37 °C and 5% CO_2_, 20 μL CellTiter 96^®^ AQueous One Solution Reagent was added per well and plates were incubated for 1.5 to 2 h at 37 °C and 5% CO_2_, prior to recording absorbance at 492 nm using a FLUOstar OPTIMA 96-well plate reader (BMG LABTECH, Offenburg, Germany). Absorbance recorded in each well was related to the average absorbance of nontreated control wells to calculate the percentage of cell viability. Datapoints are means of triplicates with SEM. Sigmoidal dose response curves were fitted and median cytotoxic concentration (CC50) values were calculated with GraphPad Prism 8.0.0 using a bottom constraint of 0 and the formula Y= Top/(1+10^((LogCC50-X)*HillSlope)) as further specified in Figures S5 and S8. To rule out cytotoxic effects at the concentrations selected based on cell viability assays in the presence of viral infection, culture wells in antiviral assays were manually inspected in the light microscope.

## Supporting information

Supplementary experimental data and descriptions

## List of abbreviations

(API): Active pharmaceutical ingredients
(CC50): median cytotoxic concentration
(COVID-19): coronavirus disease 2019
(DMSO): dimethylsulfoxide
(EC50): median effective concentration
(FBS): fetal bovine serum
(FDA): Food and Drug Administration
(IF): immunofluorescence
(PBS): phosphate buffered saline
(SD): standard deviation
(SEM): standard error of the mean
(SI): selectivity index
(SARS-CoV-2): severe acute respiratory syndrome coronavirus 2

## Acknowledgements

We thank the Max Planck Society for financial support. This work was supported by a PhD stipend from the China Scholarship Council (Y.Z.) and a grant from the Danish Agency for Science and Higher Education (J.B.). We thank Dr. Bjarne Ø. Lindhardt (Copenhagen University Hospital, Hvidovre) and Prof. Carsten Geisler (University of Copenhagen) for support from Hvidovre Hospital and the University of Copenhagen. We thank Lotte Mikkelsen, Anna-Louise Sørensen, and Pia Pedersen (Copenhagen University Hospital, Hvidovre) for laboratory assistance. We thank Dr. Christoph Rademacher and Felix Fuchsberger for helpful conversations. We thank ArtemiLife Inc. for providing the *A. annua* plant material. We thank Prof. Jean Dubuisson and Dr. Sandrine Belouzard for providing VeroE6 cells.

## Author Contributions

K.G. and P.H.S. conceived this project. K.G., Y.Z., S.R., J.B., K.O, J.M.G. and P.H.S. designed the experiments. K.G. carried out the extractions and sample preparations. K.O. and J.T. conducted the plaque-reduction assays. Y.Z., S.R., L.P., U.F., S.F. and A.O contributed to isolation of SARS-CoV-2/human/Denmark/DK-AHH1/2020 and established experimental systems. Y.Z. conducted the high-throughput antiviral assays. All authors analyzed the data. All authors contributed to and discussed the manuscript.

## Author Information

Correspondence relating to plaque-reduction assays should be addressed to K.O. Correspondance relating to high-throughput antiviral assays and cell viability assays should be addressed to J.M.G. Correspondence concerning extractions and pure compounds should be addressed to P.H.S.

