## Supplementary experimental data and descriptions for "*In vitro* efficacy of Artemisinin-based treatments against SARS-CoV-2"

### SUPPLEMENTARY INFORMATION

#### 1 Contents

1. Reagents and Materials ........ S2

2. Extraction of *A. annua* leaves ........ S3

General procedure A: Extraction using absolute ethanol ........ S3

General procedure B: extraction using distilled water ........ S3

**Extraction of *A. annua*** ........ S4

**Sample preparation** ........ S4

3. Artemisinin Stability Tests ........ S5

3.1 Thermal stability of Artemisinin ........ S5

3.2 Stability of Artemisinin in Water in the Presence of Additives ........ S5

4. Efficacy of *A. annua* extracts and artemisinin against SARS-CoV-2 in a plaque-reduction assay in VeroE6 cells .......... S8

5. Effect of artemisinin-based treatment and diluents on SARS-CoV-2 infection and cell viability *in vitro* using high-throughput assays .......... S10

5.1 High-throughput antiviral assays in VeroE6 cells .......... S10

5.2 Influence of artemisinin-based treatment on VeroE6 cell viability .......... S12

5.3 Influence of diluents and additives on SARS-CoV-2 infection and cell viability in VeroE6 cells .......... S14

5.4 Influence of artemisinin-based treatment on Huh7.5 cell viability .......... S16

5.5 Influence of diluents on SARS-CoV-2 infection and cell viability in Huh7.5 cells .......... S17

6 References .......... S17

#### 1. Reagents and Materials

Solvents were obtained from commercial suppliers and used without further purification. Dried *Artemisia annua* leaves were obtained from ArtemiLife Inc. All crops were grown and harvested in 2019 and air-dried in partial sunlight. Leaves were stripped, ground to pieces approximately ≤ 1 mm and stored in nylon bags at room temperature and used without further manipulation. Artemisinin was previously prepared and purified by crystallization using published protocols.[S1] Crystals were ground prior to use. Artesunate and artemether were purchased from TCI, stored at room temperature, and used without further purification. Filter paper used was Rotilabo type 113A, diameter 240 mm, obtained from Carl Roth.

Ground coffee used in extractions was preground 100% Arabica beans from Café Intención. For the artemisinin stability experiments, coffee was prepared using a Rowenta CT 3818 Milano coffee maker using size 4 coffee filters (Melitta brand) as follows: 100 g of ground coffee (Café Intención) was added to the cone coffee filter in the machine and 1,350 mL of water used to make the coffee using the single setting. Coffee was then transferred to a thermos, stored at room temperature, and used without further modification.

#### 2. Extraction of *A. annua* leaves

##### General procedure A: Extraction using absolute ethanol

A magnetic stir bar (6 cm) and absolute ethanol (250 mL, VWR) were added to an Erlenmeyer flask (500 mL). The stir bar was rotated at 550 rpm using a Heidolph MR HEI-END heated stir plate. The solvent was heated to 50 ˚C using the hot plate. Dried leaf material (50 g, Figure S1) was added to the ethanol using a powder funnel and allowed to stir for 200 minutes. The samples were then removed from the stir plate and vacuum filtered using a fritted filter with filter paper. The solid material was washed with room temperature ethanol until the liquid exiting the filter was clear. The solution was then dried using a rotary evaporator and further dried for at least two hours on high vacuum.

##### General procedure B: extraction using distilled water

A magnetic stir bar (6 cm) and distilled water (250 mL, VWR) were added to an Erlenmeyer flask (500 mL). The stir bar was rotated at 550 rpm using a Heidolph MR HEI-END heated stir plate. The solvent was heated to 50 ˚C using the hot plate. Dried leaf material (25 g) was added to the temperature-stable water using a powder funnel and allowed to stir for 200 minutes. Note: half the mass of plant material was used as compared to General Procedure A due to the increased absorbance of the water into the plant material. Using 125 mL (desired ratio) or 200 mL of water resulted in a thick sludge as opposed to a freely flowing heterogeneous solution.

The samples were then removed from the stir plate and vacuum filtered using a fritted filter with filter paper. The solid material was washed with room temperature water until the liquid exiting the filter was clear. The solution was then dried using a rotary evaporator and further dried for at least 2 hours on high vacuum.

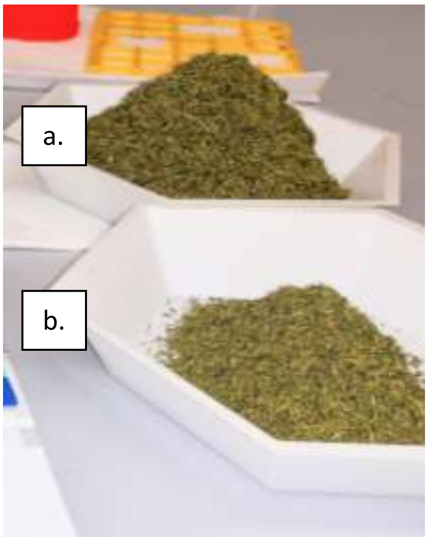


**Figure S1:** Dried leaves of *Artemisia annua*. a) Plant material (50 g). b) Plant material (4 g). Seedlings were planted in May and plants harvested in September 2019.

#### Extraction of *A. annua*

Four extracts were prepared following General Procedure A, looking at the influence of preground coffee on extraction, while one extract was prepared following General Procedure B.

***A. annua* ethanolic extract**: *A. annua* (50 g) from Lancaster, KY was added to 250 mL of 100% ethanol at 50 ˚C, stirred by magnetic stir bar. After 200 min, the plant material was removed by vacuum filtration (using a glass fritted filter and filter paper) and washed with 100% ethanol. Ethanol was removed from the solution using a rotary evaporator with water bath at 40 ˚C. The flask was further dried for three hours under high vacuum. The dried material (7.404 g) was a dark green, sticky material. The flask was covered with aluminum foil and stored at -30 ˚C prior to sample preparation.

***A. annua* + coffee ethanolic extract**: *A. annua* (50 g) from Lancaster, KY and preground coffee beans (50 g) were added to ethanol (350 mL) at 50 ˚C and stirred for 200 min. More ethanol was needed due to the viscosity of the coffee/*A. annua* mixture. Then, the plant material was removed by vacuum filtration using a glass fritted filter and filter paper and washed with ethanol. Ethanol was removed from the solution using a rotary evaporator with the water bath heated to 40 ˚C. The flask was further dried for three hours under high vacuum. The dried material (11.612 g) was a very dark green, oily material on the bottom of the flask and a dark green, sticky solid on the sides. The flask was covered with aluminum foil and stored at -30 ˚C prior to sample preparation.

**Coffee ethanolic extract (for reference).** Preground coffee (50 g) was added to ethanol (250 mL of 100%) at 50 ˚C, and stirred for 200 min, before the plant material was removed by vacuum filtration (using glass fritted filter and filter paper) and washed with ethanol. Ethanol was removed using a rotary evaporator with water bath at 40 ˚C. The flask was further dried for three hours under high vacuum. The dried material (6.838 g) appeared to be two different materials, a transparent light brown liquid and a dark brown solid. The flask was covered with aluminum foil and stored at -30 ˚C prior to sample preparation.

***A. annua* aqueaous extract**: *A. annua* (25 g) from Lancaster, KY was added to 250 mL of distilled water at 50 ˚C, stirred by magnetic stir bar. After 200 min, the plant material was removed by vacuum filtration (using a glass fritted filter and filter paper) and washed with distilled water. Water was removed from the solution using a rotary evaporator with water bath at 40 ˚C. The flask was further dried for three hours under high vacuum. The dried material (6.915 g) was a dry, green-brown material. The flask was covered with aluminum foil and stored at -30 ˚C prior to sample preparation.

#### Sample preparation

Seven samples were prepared for antiviral assays.

**Sample 1 – *A. annua* ethanolic extract.** *A.annua* ethanolic extract was dissolved in DMSO. Light heating was required for maximal solvation – trace amounts of solid material could be observed. The sample was filtered through a syringe filter (Chromafil® xtra RC0.45). The sample was stored at -20 °C until use.

**Sample 2 – *A. annua* + coffee ethanolic extract.** *A. annua* +coffee ethanolic extract was dissolved in DMSO. Light heating was required for maximal solvation as trace amounts of solid material were observed. The sample was filtered through a syringe filter (Chromafil® xtra RC0.45). The sample was stored at -20 °C until use.

**Sample 3 – *A. annua* aqueous extract.** *A. annua* aqueous extract was dissolved in DMSO:$H_2O$ (3:1). Light heating was required for maximal solvation – trace amounts of solid material could be observed. The sample was filtered through a syringe filter (Chromafil® xtra RC0.45). The sample was stored at -20 °C until use.

**Sample 4 – Artemisinin.** Artemisinin was dissolved in DMSO. Full solvation was obtained with light heating. The sample was stored at -20 °C until use.

**Sample 5 – Artesunate.** Artesunate was dissolved in DMSO. Full solvation was obtained with light heating. The sample was stored at -20 °C until use.

**Sample 6 – Artemether.** Artemether was dissolved in DMSO. Full solvation was obtained with light heating. The sample was stored at -20 °C until use.

**Sample 7 – Coffee ethanolic extract.** Coffee ethanolic extract was dissolved in DMSO. Light heating was required for maximal solvation as trace amounts of solid material were observed. The sample was filtered through a syringe filter (Chromafil® xtra RC0.45). The sample was stored at -20 °C until use.

#### 3. Artemisinin Stability Tests

##### 3.1 Thermal stability of Artemisinin

Artemisinin crystals were ground to a white powder using a glass vial. In duplicate, artemisinin (30 mg) was placed in an open glass vial (5 mL) and placed into a drying oven held at either 100 ˚C, 85 ˚C, 73 ˚C, or 66 ˚C for 15 minutes. No color change was observed. All samples were dissolved in deuterated chloroform ($CDCl_3$) containing 0.5 equivalents of an internal standard (1,2,4,5-tetramethylbenzene). No degradation of artemisinin was observed by $^1$H NMR.

##### 3.2 Stability of Artemisinin in Water in the Presence of Additives

The stability of artemisinin in aqueous solutions was tested in the presence of reductants in tap water and black coffee. Two reductants were tested, glucose (1 equivalent) and Fe(II)$Cl_2$•4$H_2O$ (0.5

equivalents). All mixtures were heated to 93 ˚C for 90 minutes in 10 mL glass vials open to the atmosphere. Halfway through heating tap water (1-2 mL) was added to compensate for evaporation. Following heating, ethyl acetate (2 mL) was added, and the biphasic solution was dried using a rotary evaporator with a water bath at 40 ˚C. The flask was further dried for three hours under high vacuum. All samples were dissolved in $CDCl_3$ containing 0.5 equivalents of an internal standard (1,2,4,5-tetramethylbenzene) and $^1$H NMR performed.

*Aqueous Stability 1:* Ground artemisinin (30.8 mg) was added to tap water (5 mL). The mixture was heated for 90 min prior to cooling, ethyl acetate addition, and evaporation to yield 36.4 mg of a white solid. $^1$H NMR revealed no degradation of artemisinin.

*Aqueous Stability 2:* Ground artemisinin (30.8 mg) and glucose (18.2 mg, one equivalent) were added to tap water (5 mL). The mixture was heated for 90 min prior to cooling, ethyl acetate addition, and evaporation to yield 64.1 mg of a yellowish oil/plaque with white spots. $^1$H NMR revealed 32% loss of artemisinin. No degradation products were identified.

*Aqueous Stability 3:* Ground artemisinin (29.5 mg) and $Fe(II)Cl_2{\bullet}4H_2O$ (11.4 mg, 0.5 equivalents) were added to tap water (5 mL). The mixture was heated for 90 min prior to cooling, ethyl acetate addition, and evaporation to yield 92.6 mg of a pink/orange solid. $^1$H NMR revealed complete loss of artemisinin. No degradation products were identified.

*Aqueous Stability 4:* Ground artemisinin (30.4 mg), glucose (19.4 mg, 1 equivalent) and $Fe(II)Cl_2{\bullet}4H_2O$ (11.4 mg, 0.5 equivalents) were added to tap water (5 mL). The mixture was heated for 90 min prior to cooling, ethyl acetate addition, and evaporation to yield 67.3 mg of a black solid. $^1$H NMR revealed complete loss of artemisinin. No degradation products were identified.

*Aqueous Stability 5:* Ground artemisinin (30.6 mg) was added to premade black coffee (5 mL). The mixture was heated for 90 min prior to cooling, ethyl acetate addition, and evaporation to yield 175 mg of brown/black solid. $^1$H NMR revealed 17% degradation of artemisinin. No degradation products were identified.

*Aqueous Stability 6:* Ground artemisinin (30.5 mg) and glucose (19.6 mg, one equivalent) were added to black coffee (5 mL). The mixture was heated for 90 min prior to cooling, ethyl acetate addition, and evaporation to yield a brown/black solid (302 mg). $^1$H NMR revealed 19% loss of artemisinin. No degradation products were identified.

*Aqueous Stability 7:* Ground artemisinin (30.2 mg) and $Fe(II)Cl_2{\bullet}4H_2O$ (12.4 mg, 0.5 equivalents) were added to black coffee (5 mL). The mixture was heated for 90 min prior to cooling, ethyl acetate addition, and evaporation to yield 192 mg of a black solid. $^1$H NMR revealed complete loss of artemisinin. No degradation products were identified.

*Aqueous Stability 8:* Ground artemisinin (30.3 mg), glucose (18.5 mg, 1 equivalent) and Fe(II)$Cl_2$•4$H_2O$ (10 mg, 0.5 equivalents) were added to black coffee (5 mL). The mixture was heated for 90 min prior to cooling, ethyl acetate addition, and evaporation to yield 198 mg of a black solid. $^1$H NMR revealed complete loss of artemisinin. No degradation products were identified.

#### 4. Efficacy of *A. annua* extracts and artemisinin against SARS-CoV-2 in a plaque-reduction assay in VeroE6 cells

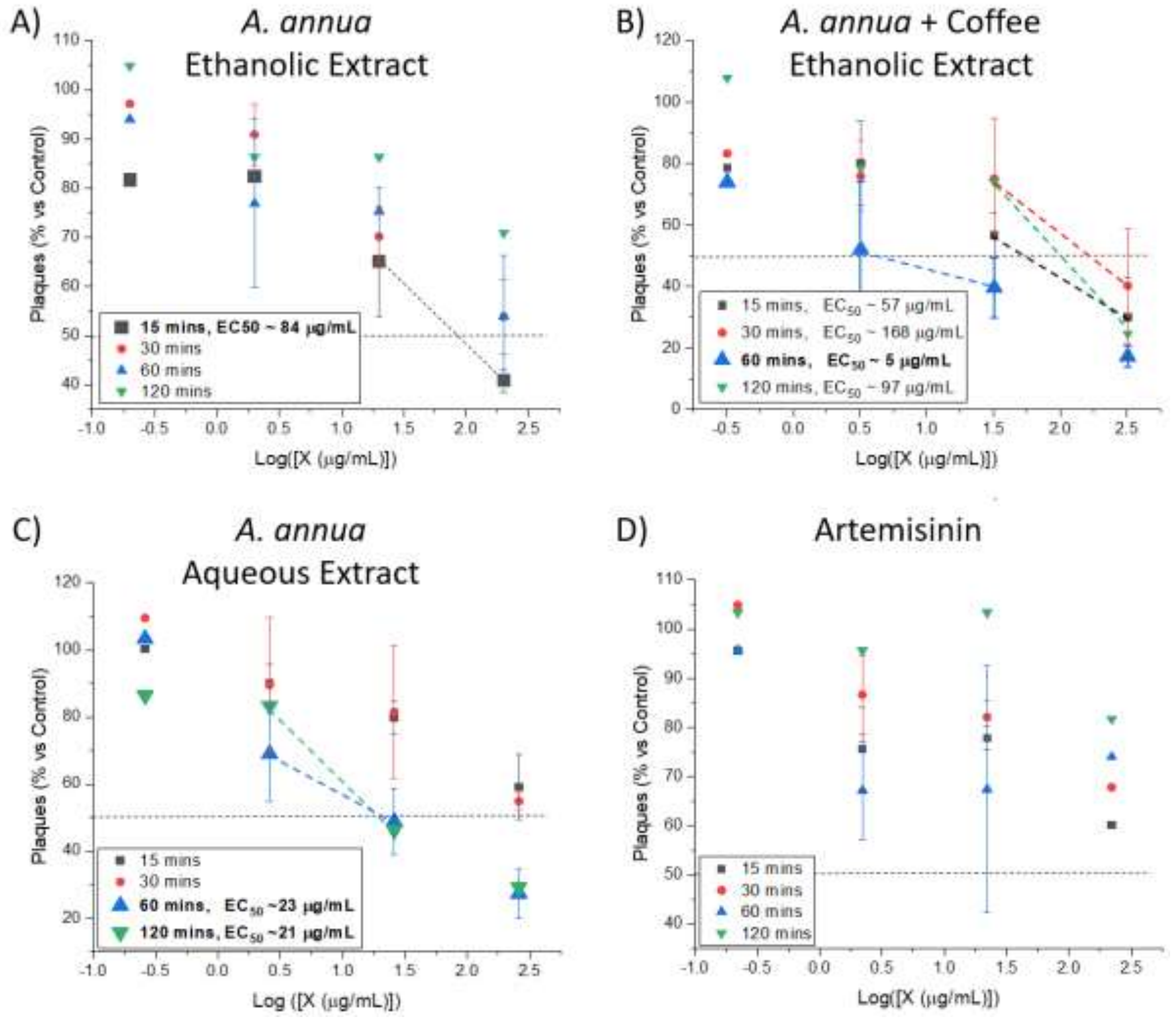


**Figure S2. Antiviral activity of *A. annua* extracts and artemisinin against SARS-CoV-2 using a plaque-reduction assay in VeroE6 cells.** VeroE6 cells were treated with the extracts or pure artemisinin for either 15, 30, 60, or 120 minutes prior to infection with isolate SARS-CoV-2M at ~130 plaque-forming units per well. DMSO was used as a negative control and plaque numbers were determined by manual counting following indirect immunofluorescence using a mixture of antibodies to SARS-CoV Nuclear protein N or following staining with crystal violet. $EC_{50}$ values were calculated using the equation $EC_{50}$ = 10^x, where x is obtained by solving the equation of the line between the two points on either side of 50% inhibition (when available) using the formula y = mx+b, where y = 50. For data where 50% viral inhibition was not achieved, no $EC_{50}$ value was calculated. The line equations for each $EC_{50}$ value calculated are given as follows. (A) *A. annua* ethanolic extract: (15 min)

y = -24.323x + 96.844; (B) *A. annua* + coffee ethanolic extract: (15 min) y = -26.577x + 96.658; (30 min) y = -34.824x + 127.52; (60 min) y = -12.458x + 58.38; (120 min) y = -49.357x + 148.33; (C) *A. annua* aqueous extract: (60 min) y = -20.23x + 77.449; (120 min) y = -37.018x + 98.652; (D) artemisinin: 50% viral inhibition was not achieved. Datapoints are based on two replicates from two independent assays; error bars reflecting standard deviations (SD) are shown. For datapoints without error bars, only one replicate was available.

### 5. Effect of artemisinin-based treatment and diluents on SARS-CoV-2 infection and cell viability *in vitro* using high-throughput assays

#### 5.1 High-throughput antiviral assays in VeroE6 cells

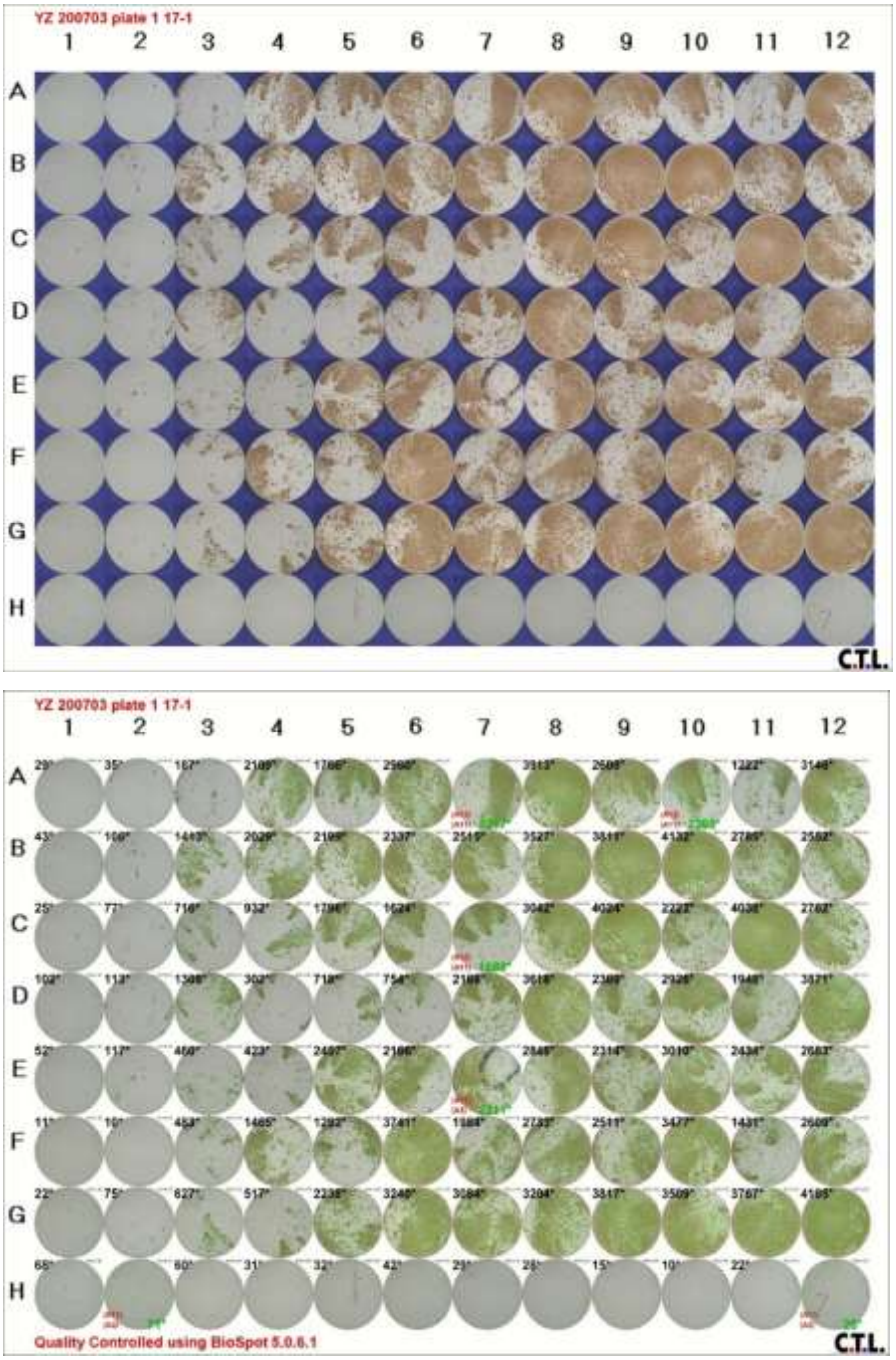


**Figure S3. Representative 96-well images from high-throughput antiviral assays in VeroE6 cells.** A treatment high-throughput antiviral assay was carried out in VeroE6 cells as described in Materials and Methods using the *A. annua* ethanolic extract. An image of the complete 96-well plate is shown prior to (top image) and following (bottom image) automated counting of single SARS-CoV-2 spike glycoprotein positive cells.

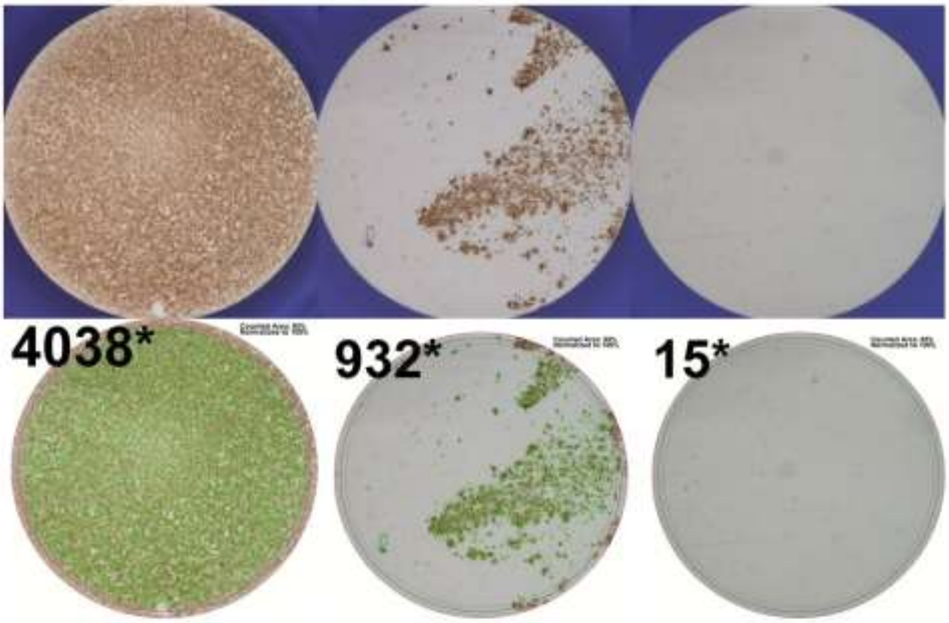


**Figure S4. Representative single-well images from high-throughput antiviral assays in VeroE6 cells.** A treatment high-throughput antiviral assay was carried out in VeroE6 cells as described in Materials and Methods using the *A. annua* ethanolic extract. Representative images of single wells showing different degrees of SARS-CoV-2 infection are shown prior to (first row) and following (second row) counting of single SARS-CoV-2 spike glycoprotein positive cells. Single-well images are derived from Figure S3 wells C11, C4, and H9.

#### 5.2 Influence of artemisinin-based treatment on VeroE6 cell viability

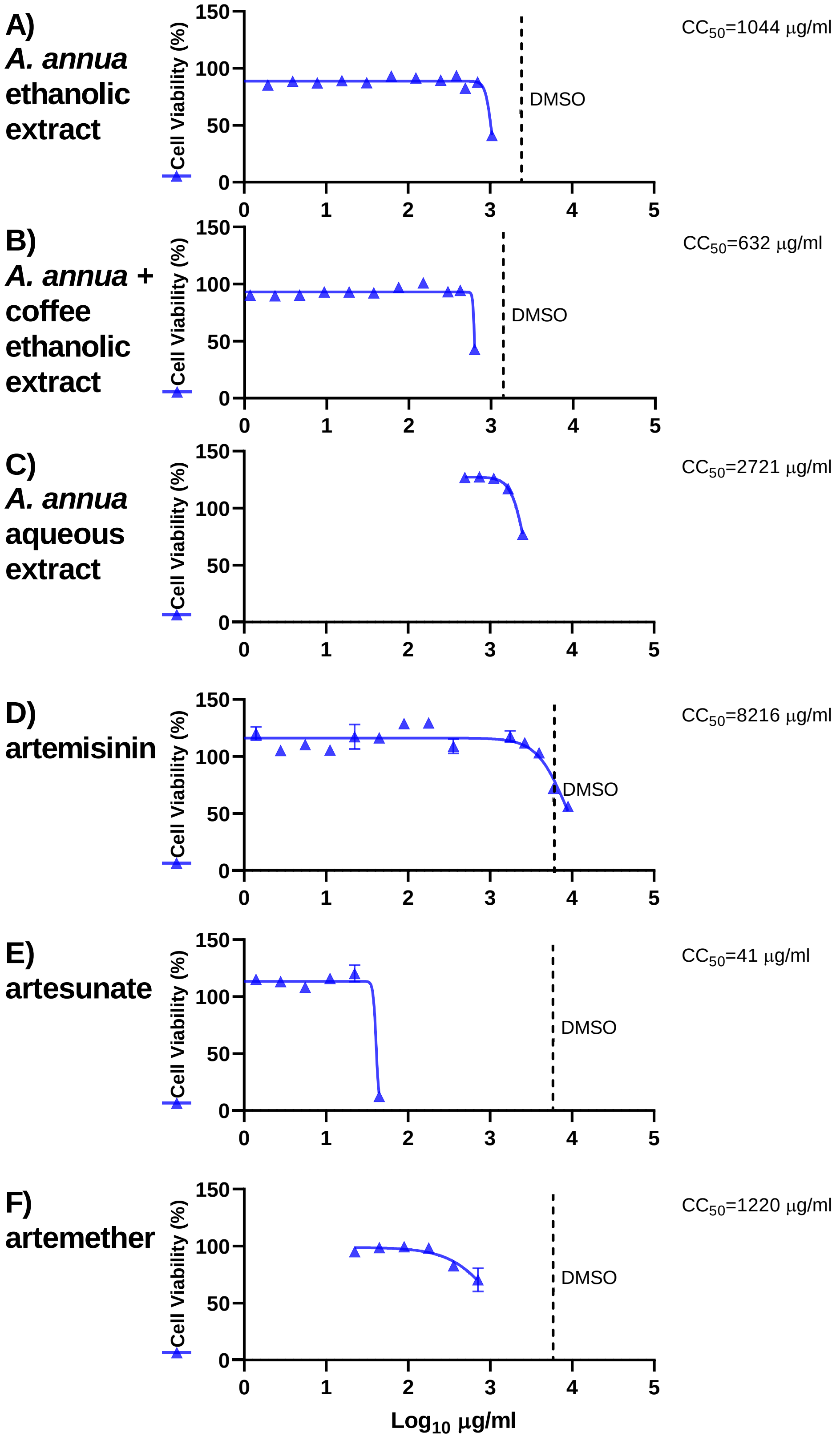

**Figure S5. Calculation of CC50 based on cell viability assays in VeroE6 cells.** Cell viability assays were carried out as described in Materials and Methods using the *A. annua* ethanolic extract (A), *A. annua* + coffee ethanolic extract (B), *A. annua* aqueous extract (C), or compounds artemisinin (D), artesunate (E), and artemether (F). All datapoints are shown in Figures 2 and 3 of the manuscript. To facilitate fitting of sigmoidal dose response curves and calculation of CC50 values, of the datapoints providing the lower plateau of the curve, only the datapoint at the lowest concentration was included in the analysis. Sigmoidal dose response curves were fitted in GraphPad Prism 8.0.0 using a bottom constraint of 0 and the formula Y= Top/(1+10^((LogEC50-X)*HillSlope)).

#### 5.3 Influence of diluents and additives on SARS-CoV-2 infection and cell viability in VeroE6 cells

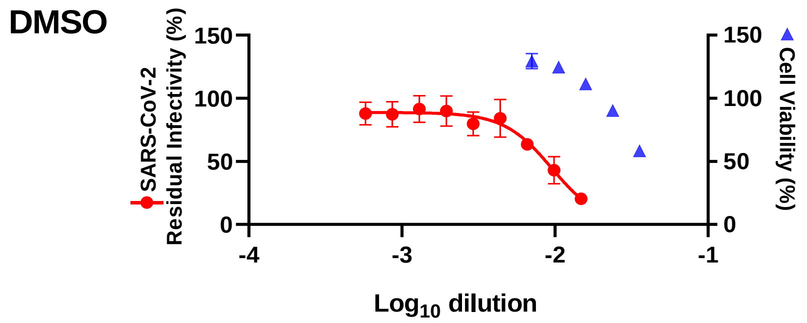


**Figure S6. Effect of DMSO on SARS-CoV-2 and cell viability in VeroE6 cells.** VeroE6 cells seeded the previous day in 96-well plates were infected with SARS-CoV-2 and after 1 hour incubation treated with the specified dilutions of DMSO. After a 2-day incubation, infected cells were visualized by immunostaining for SARS-CoV-2 spike glycoprotein and counted automatically as described in Materials and Methods. % residual infectivity for individual wells was calculated by relating counts of infected treated wells to the mean count of 14 infected nontreated control wells. Datapoints (red dots) are means of seven replicates with standard error of the means (SEM). Sigmoidal dose response curve (red line) was fitted in GraphPad Prism as described in Materials and Methods. % Cell viability was determined in replicate assays without infection with SARS-CoV-2 as described in Materials and Methods. Datapoints (blue triangles) are means of three replicates with SEM.

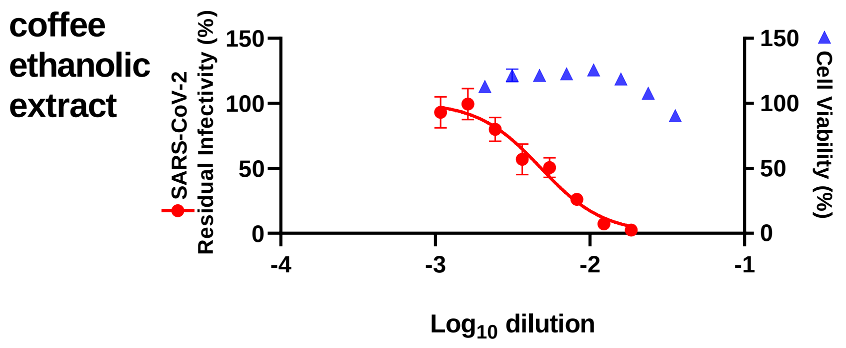


**Figure S7. Effect of coffee ethanolic extract on cell viability and SARS-CoV-2 in VeroE6 cells.** VeroE6 cells seeded the previous day in 96-well plates were infected with SARS-CoV-2 and after 1 hour incubation treated with the specified dilutions of coffee ethanolic extract. After a 2-day incubation, infected cells were visualized by immunostaining for SARS-CoV-2 spike glycoprotein and counted automatically as described in Materials and Methods. % residual infectivity for individual wells was calculated by relating counts of infected treated wells to the mean count of 14 infected nontreated control wells. Datapoints (red dots) are means of seven replicates with SEM. Sigmoidal dose response curve (red line) was fitted in GraphPad Prism as described in Materials and Methods. % Cell viability was determined in replicate assays without infection with SARS-CoV-2 as described in Materials and Methods. Datapoints (blue triangles) are means of 3 replicates with SEM.

#### 5.4 Influence of artemisinin-based treatment on Huh7.5 cell viability

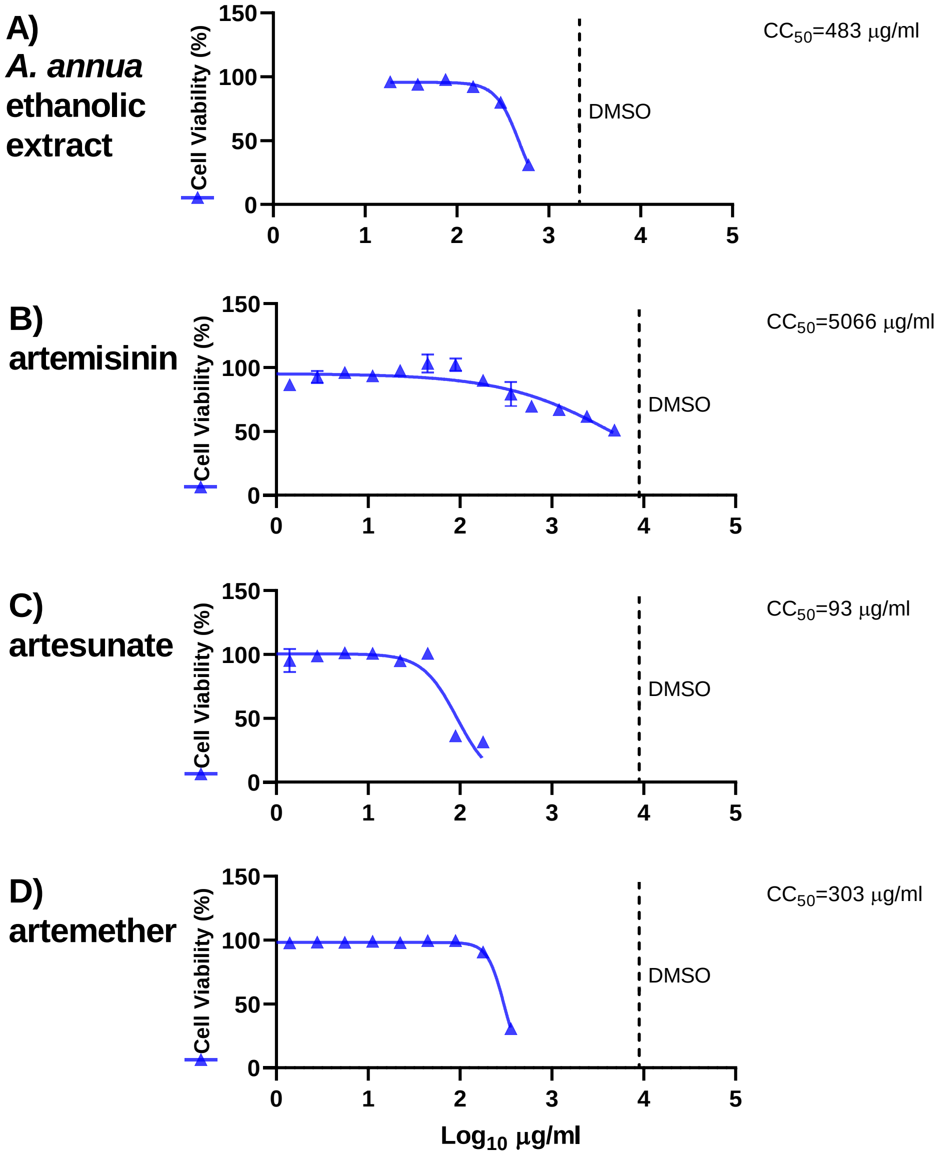


**Figure S8. Calculation of CC50 based on cell viability assays in Huh7.5 cells.** Cell viability assays were carried out as described above using the *A. annua* ethanolic extract (A), or compounds artemisinin (B), artesunate (C), and artemether (D). All datapoints are shown in Figure 4 of the manuscript. To facilitate fitting of sigmoidal dose response curves and calculation of CC50 values, of the datapoints providing the lower plateau of the curve, only the datapoint at the lowest concentration was included in the analysis. Sigmoidal dose response curves were fitted in GraphPad Prism 8.0.0 using a bottom constraint of 0 and the formula Y= Top/(1+10^((LogEC50-X)*HillSlope)).

#### 5.5 Influence of diluents on SARS-CoV-2 infection and cell viability in Huh7.5 cells

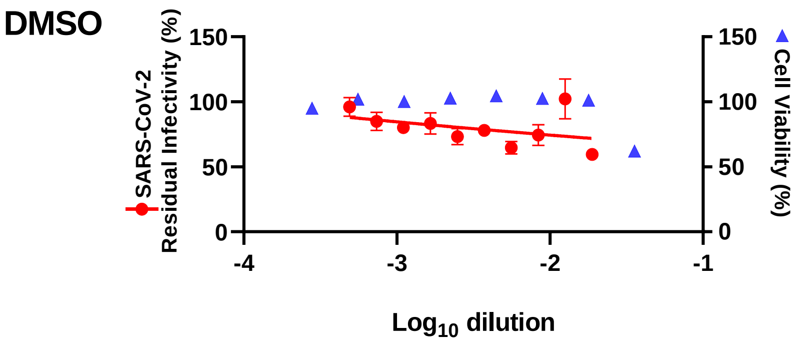


**Figure S9. Effect of DMSO on cell viability and SARS-CoV-2 in Huh7.5 cells.** Huh7.5 cells seeded the previous day in 96-well plates were infected with SARS-CoV-2 and directly treated with the specified dilutions of DMSO. After a 3-day incubation, infected cells were visualized by immunostaining for SARS-CoV-2 spike glycoprotein and counted automatically as described in Materials and Methods. % residual infectivity for individual wells was calculated by relating counts of infected treated wells to the mean count of 14 infected nontreated control wells. Datapoints (red dots) are means of seven replicates with SEM. Sigmoidal dose response curve (red line) was fitted in GraphPad Prism as described in Materials and Methods. % Cell viability was determined in replicate assays without infection with SARS-CoV-2 as described in Materials and Methods. Datapoints (blue triangles) are means of three replicates with SEM.
